# Persistent effects of the orexin-1 receptor antagonist SB-334867 on motivation for the fast acting opioid remifentanil

**DOI:** 10.1101/521633

**Authors:** Aida Mohammadkhani, Morgan H. James, Gary Aston-Jones

**Author notes:** Submitting author: Aida Mohammadkhani. Corresponding author: Gary Aston-Jones, Ph.D. Brain Health Institute Rutgers University 683 Hoes Lane West, R. 259 Piscataway NJ 08854, E. For submission to the special issue: *‘Orexin/hypocretin receptor antagonists for the treatment of addiction and related psychiatric disease: What are the steps from here?’ (Eds. Aston-Jones and James)*.

## Abstract

The orexin (hypocretin) system is important for reward-seeking behavior. The orexin-1 receptor (Ox1R) antagonist SB334867 (SB) reduces seeking of food and drug reward under conditions of high motivation. There is some evidence that the effects of systemic SB on reward seeking persist beyond the pharmacological availability of the drug, however the time course of these effects is not well characterized, nor is it known whether similar persistent effects are observed following intraparenchymal injections. Here, we used a behavioral economics paradigm, which allows for repeated testing of drug motivation across consecutive days, to examine the persistent effects of acute systemic and local treatment with SB on motivation for the short-acting opioid remifentanil. Systemic injections of SB immediately prior to behavioral testing reduced motivation for remifentanil; this effect was sustained on a subsequent test at 24h, but not on a third test at 48h. When injected locally into caudal ventral pallidum (cVP) the effects of SB were more persistent, with reduced motivation observed for up to 48h. We next made SB injections into cVP 24h prior to behavioral testing; this produced persistent effects that persisted for at least 72h post-treatment. Cued reinstatement of extinguished remifentanil seeking was also attenuated by pretreatment with SB 24h prior. These data indicate that the effects of SB on opioid seeking behavior persist beyond the bioavailability of the compound. These observations might have important ramifications for the future clinical use of orexin receptor antagonists for the treatment of addiction.

## 1. Introduction

Orexin peptides A and B (also known as hypocretin 1 and 2) are produced by hypothalamic neurons (de Lecea et al., 1998; Sakurai et al., 1998b) and are important for motivation for food and drugs of abuse (Barson, 2018; Borgland et al., 2009; Brodnik et al., 2018; Choi et al., 2010; Harris et al., 2005; James et al., 2017; Mahler et al., 2014a; Walker and Lawrence, 2017; Yeoh et al., 2018). Orexinergic fibers project widely throughout the brain (Baldo et al., 2003; Peyron et al., 1998) to release orexin peptides A and B which act at two G-protein coupled receptors, orexin receptor 1 and 2 (Ox1R and Ox2R) (de Lecea et al., 1998; Hervieu et al., 2001; Marcus et al., 2001; Sakurai et al., 1998a). Studies generally support a dichotomy of function for Ox1R versus Ox2R signaling, whereby signaling at Ox1R is important for motivation and reward, and signaling at Ox2R is more involved in arousal processes (Brodnik et al., 2018). Accordingly, a significant body of literature shows that administration of Ox1R antagonists, either systemically or locally into reward regions, attenuates a broad range of drug-seeking behaviors (Bentzley and Aston-Jones, 2015; Espana et al., 2010; Harris et al., 2005; James et al., 2011; James et al., 2018a; Jupp et al., 2011; Mahler et al., 2013; Moorman et al., 2017; Porter-Stransky et al., 2017; Schmeichel et al., 2017; Smith and Aston-Jones, 2012). Such findings have spurred speculation that orexin receptor antagonists might represent a novel and effective option for the treatment of addiction (Brodnik et al., 2018; Campbell et al., 2018; James and Aston-Jones, 2017; James et al., 2017; Khoo and Brown, 2014; Perrey and Zhang, 2018; Walker and Lawrence, 2017; Yeoh et al., 2014).

Of the Ox1R antagonists available, SB-334867 (SB) is by far the most commonly used in animal studies, with >400 publications reporting the use of this compound (Perrey and Zhang, 2018). At a dose of 30mg/kg administered systemically, SB is effective at reducing motivated seeking of all drugs tested with limited/no effect on arousal or general motor activity (James et al., 2018c; LeSage et al., 2010; Moorman and Aston-Jones, 2009; Porter-Stransky et al., 2017; Smith and Aston-Jones, 2012). At this dose, peak blood and brain levels are observed 30min post-dosing; high levels are maintained at 4h post-dosing but gradually decline to be virtually non-detectable at 8h (Ishii et al., 2005). Accordingly, studies using SB generally commence behavioral testing 15-30mins after SB administration, and testing is conducted over 1-2h when brain-SB concentrations remain high (Bentzley and Aston-Jones, 2015; Borgland et al., 2006; James et al., 2018c; Smith et al., 2009; Smith et al., 2010; Smith and Aston-Jones, 2012). In studies where SB is delivered locally into discrete brain regions, behavioral testing typically commences within 10min of SB injections (Harris et al., 2007; James et al., 2011; Mahler et al., 2013; Narita et al., 2006).

Despite the relatively short pharmacokinetic profile of SB, two studies reported that the behavioral effects of this compound on feeding behavior are maintained beyond its bioavailability. Both studies showed that the anorexic effects of SB on palatable food or chow intake were maintained at 24h post-acute dosing, an effect that was also accompanied by a significant loss of bodyweight (Haynes et al., 2000; Ishii et al., 2005). One of these studies measured feeding and weight gain at 48h and showed that both indices were normalized at this time point (Ishii et al., 2005). Despite the many studies examining the effect of SB on drug seeking, the persistence of these effects is poorly understood. To our knowledge, only one paper has addressed this question directly. Brodnick et al. (2018) recently reported that systemic injections of SB reduce cocaine self-administration for at least 21h post-delivery, indicating that the anti-drug seeking properties of SB persist beyond its availability. Currently, it is not known how long these effects persist, nor if they extend to local (intracranial) injections of SB. Moreover, it is unclear whether acute SB administration is also associated with a persistent suppression of self-administration for other drugs of abuse, including opioids. Here, we addressed these questions directly by using a behavioral economics (BE) measure of demand/motivation for the short-acting opioid remifentanil. We chose this approach because inter-individual behavior on this task is highly stable (Bentzley et al., 2014; James et al., 2018a), allowing for repeated testing of drug motivation over consecutive days. In addition, we previously reported that SB, delivered either systemically or directly into caudal ventral pallidum (cVP), suppresses motivation for remifentanil on a BE task (Mohammdkhani et al., 2018; Porter-Stransky et al., 2017); thus we could directly compare the persistence of effects across these two distinct routes of administration.

We report that systemic SB reduces motivation for remifentanil on an initial demand test (0.5h) and on a subsequent test at 24h post-treatment, but not on a third test at 48h. The effects of SB were more persistent when administered locally into cVP, as reductions in motivation for remifentanil were observed at 0, 24 and 48h post-treatment. We further show that when administered 24h prior to behavioral testing, intra-cVP injections of SB are effective at reducing drug motivation, and that these effects persist for 72h. Together our data indicate that SB delivered systemically or locally into cVP has effects on remifentanil motivation that persist beyond the pharmacological availability of the compound.

## 2. Results

### 2.1. Experiment 1

#### Systemic SB30 administration reduced demand for remifentanil up to 24h post-treatment

We reported elsewhere (Mohammdkhani et al., 2018) that acute systemic treatment with SB prior to behavioral testing reduced motivation for remifentanil (0.5h). Here, we tested the persistence of these effects by analyzing behavior in subsequent BE tests at 24h and 48h post-SB. A two-way ANOVA of α values following SB or vehicle treatment revealed a significant ‘Treatment’ x ‘Day’ interaction (F_3,54_=3.585, p=0.0195). Subsequent post-hoc analyses indicated that SB treatment was associated with significantly increased α values (decreased motivation) compared to baseline at 0.5h and 24h (baseline vs. 0.5h, p=0.0003; baseline vs. 24h, p=0.0442), but not 48h (baseline vs. 48h: p=0.9502), following SB treatment (**Fig. 1a**). Vehicle treatment had no effect on α values at any time point (**Fig. 1a**). A similar analysis of Q_0_ values (remifentanil intake at low effort) revealed a ‘Treatment’ x ‘Day’ interaction (**Fig. 1b**; F_3,54_=3.473, p=0.0222); post-hoc analyses indicated that Q_0_ values were significantly reduced at 0.5h (baseline vs. 0.5h: p=0.0013), but not at 24h or 48h (baseline vs. 24h: p>0.05; baseline vs. 48h: p>0.05; **Fig. 1b**). There was no effect of vehicle on Q_0_ values (**Fig. 1b**).

**Figure 1.**
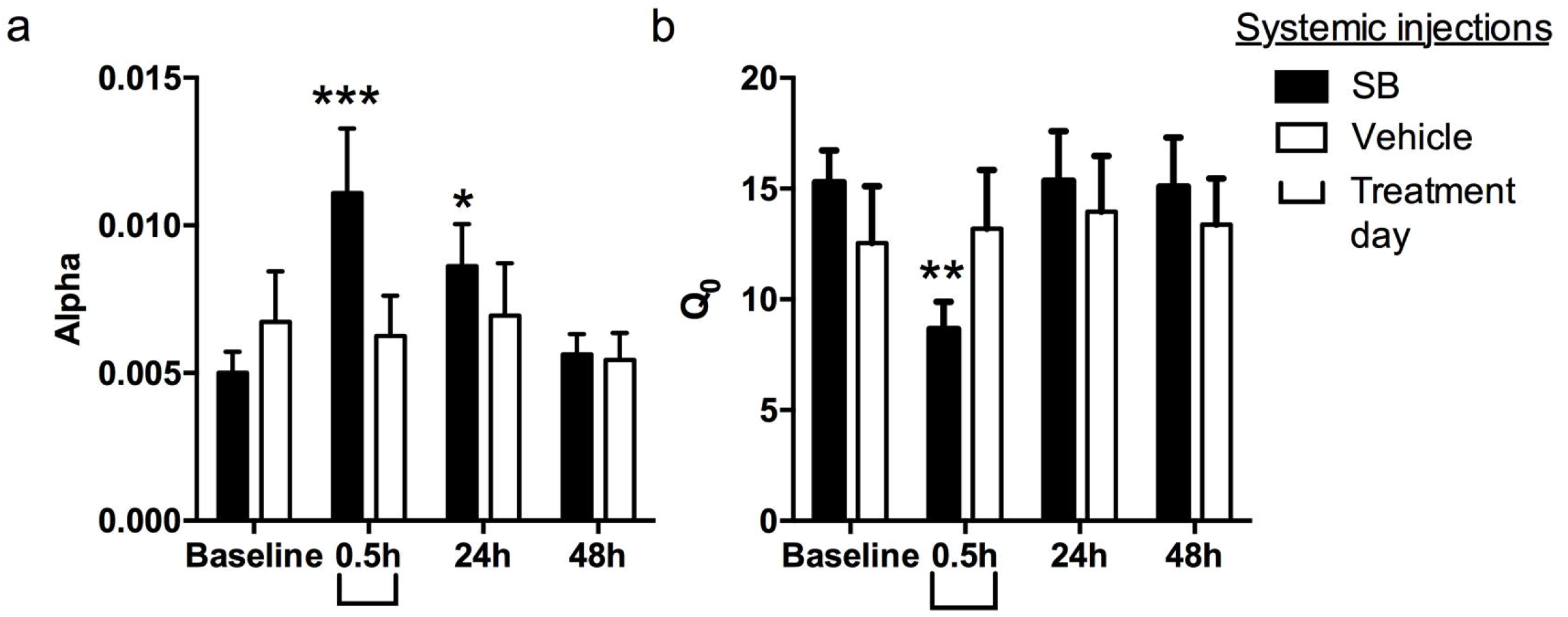
Systemic injections of SB reduced motivation for remifentanil up to 24h post-treatment. **(a)** Animals were treated systemically with SB (30mg/kg) or vehicle immediately prior to testing on a behavioral economics paradigm for remifentanil demand (0.5h). BE testing was repeated (with no further SB administration) at 24h and 48h post-treatment. Compared to baseline values, SB-treated animals exhibited a significant increase in alpha values (decreased motivation) at 0.5h and 24h post-treatment. Alpha values returned to baseline levels at 48h post-SB administration. There was no change in alpha values following vehicle treatment. **(b)** Systemic SB administration reduced remifentanil free consumption at null cost (Q_0_) at 0.5h only. Bar graphs represent mean ± standard error of mean (SEM). *p<0.05, **p<0.01, ***p<0.001 compared to baseline values.

### 2.2. Experiment 2

#### Intra-VP SB microinfusions reduced demand for remifentanil up to 48h post-treatment

We also reported that intra-cVP injections of SB immediately prior to behavioral testing reduced demand for remifentanil (Mohammadkhani et al., 2018). Here we tested the persistence of these effects by analyzing data from subsequent BE sessions at 24h and 48h post-SB intra-cVP. A repeated-measures ANOVA of α values following SB/vehicle treatment revealed a significant ‘Treatment’ x ‘Day’ interaction (Experiment 2: F_3,78_=4.13, p=0.0090). Subsequent post-hoc analyses revealed that α values were significantly increased (motivation was decreased) at 0h (baseline vs. 0h: p<0.0001), 24h (baseline vs. 24h: p<0.0001) and 48h (baseline vs. 48h: p<0.05) post-SB, reflecting a persistent decrease in motivation for remifentanil (**Fig. 2a**). There was no effect of aCSF microinjections on α values (**Fig. 2a**). A similar analysis of Q_0_ values revealed no effect of SB or aCSF at any time point (F_3, 78_=0.4549, p=0.714; **Fig. 2b**).

**Figure 2.**
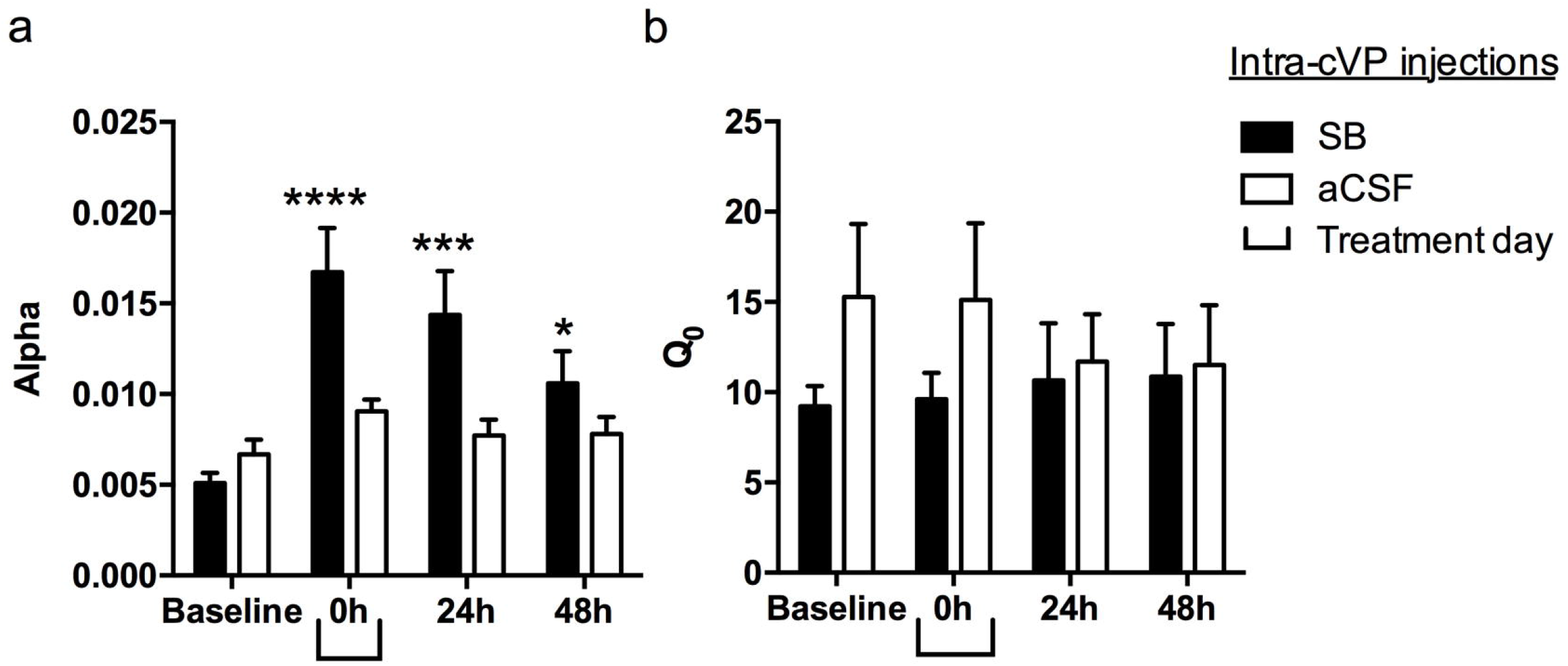
Intra-cVP infusions of SB reduced motivation for remifentanil for at least 48h post-treatment. **(a)** Animals received intra-cVP microinfusions of SB (1mM) or aCSF immediately prior to testing on a BE paradigm for remifentanil demand (0h). BE testing was repeated (with no further SB administration) at 24h and 48h post-treatment. Compared to baseline values, intra-cVP SB treatment was associated with a significant increase in alpha (decreased motivation) at 0h, 24h and 48h post-treatment. There was no effect of aCSF treatment. **(b)** There was no effect of SB on remifentanil consumption at null cost (Q_0_). Bar graphs represent mean ± standard error of mean (SEM). *p<0.05, ***p<0.001, ****p<0.0001, compared with baseline values.

### 2.3. Experiment 3

#### 2.3.1. Intra-cVP pretreatment with SB (24h prior to testing) reduced demand for remifentanil for at least 72h

A key question emerging from *Experiment 2* was whether plasticity or other changes associated with reduced remifentanil intake in the initial behavioral test following intra-cVP SB treatment contributed to reduced motivation for remifentanil in subsequent test sessions. To test this, we trained a new group of rats on the BE paradigm and microinjected SB 24h prior to the first behavioral test (**Fig. 3a**). Similar to the groups used for Experiments 1 and 2 (Mohammdkhani et al., 2018), these rats exhibited a robust preference for the active versus the inactive lever over the final 6d of self-administration training (**Fig. 3b**; RM ANOVA; F_10,131_= 13.21, p<0.001). Active lever responding increased across training and reached a plateau over the final 3 sessions (**Fig. 3b**). The total number of remifentanil infusions earned over the training period was similar to that of animals included in Experiments 1 and 2 (data shown in Mohammadkhani et al., 2018). Responding on the BE task resulted in accurate demand curve fitting; a representative curve is presented in **Fig. 3c**. Baseline α and Q_0_ values were similar to those from animals in Experiments 1 and 2 (also reported in Mohammadkhani et al., 2018). Similar to previous reports (Mohammdkhani et al., 2018; Porter-Stransky et al., 2017), α and Q_0_ values were negatively correlated (**Fig. 3d**; r=−0.6243, p=0.0401).

**Figure 3.**
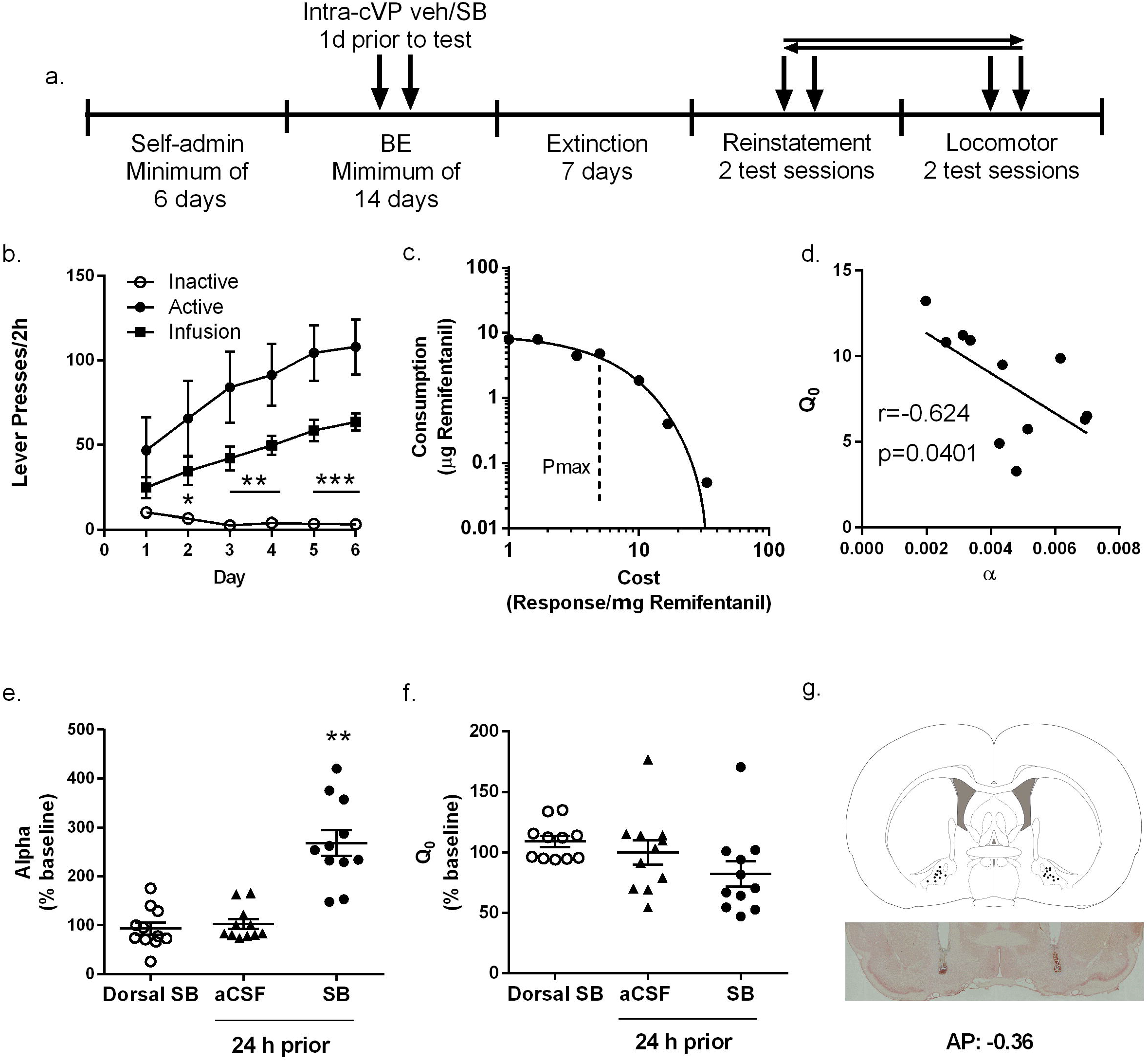
Intra-cVP pretreatment with SB 24h prior to testing reduced motivation for remifentanil. **(a)** Schematic of behavioral training and testing in Experiment 3 (Figs. 3-5), where intra-cVP microinjections of SB or aCSF were made 24h prior to BE testing sessions. **(b)** Mean behavioral performance during remifentanil self-administration training (n=11). Data show average number of infusions, active and inactive lever presses during the last 6 days of FR1 sessions. Statistical symbols represent comparisons between active and inactive lever responses. **(c)** A representative demand curve of a single animal during a single remifentanil BE session. **(d)** Remifentanil intake at low cost (Q_0_) and demand elasticity (α) are negatively correlated. **(e)** When administered 24h prior to demand testing, intra-cVP SB injections significantly increased demand elasticity (α; lowered motivation) for remifentanil, compared to aCSF or microinfusion of SB dorsal to cVP. Statistical symbols represent comparison with aCSF control group. **(f)** Pretreatment with SB had no effect on consumption of remifentanil at null cost (Q_0_). **(g)** Representative schematic of VP (top panel; adapted from Paxinos & Watson brain atlas) depicting injection sites from animals in Experiment 3, based on inspection of brain tissue following neutral red stain (representative section shown in lower panel; frontal section, midline at center). **p<0.01.

After baseline BE measures were obtained, rats were pretreated 24h prior to testing on subsequent BE sessions. Repeated measures ANOVA revealed that microinfusion of SB into cVP 24h prior to BE testing significantly increased demand elasticity compared to microinfusion of aCSF or dorsal infusions of SB (**Fig. 3e**; F_2,32_=28.92, p=0.0001). Importantly, the effect of SB pretreatment on α values was not due to upward diffusion of SB along the cannulae tract, as α values following dorsal SB infusions were no different to vehicle treatment (Fig 3e; p=0.58). Despite an overall significant effect of treatment on remifentanil free consumption (Q_0_; **Fig. 3f**; F_2,32_=4.268, p=0.0394), post-hoc tests failed to find significant differences between groups (p’s>0.05).

To test the persistence of these effects, we performed a two-way ANOVA of α values from subsequent daily BE tests. This revealed a significant ‘Treatment’ x ‘Day’ interaction (**Fig. 4a**: F_4,80_=7.032, p<0.0001), with subsequent posthoc tests indicating that SB pretreatment was associated with significantly higher α values (lower motivation) at 24h (p<0.0001), 48h (p=0.0002) and 72h (p=0.0166) post-treatment (**Fig. 4a**). There was no effect of vehicle treatment on Q_0_ values at any time point (**Fig. 4b**; F_4, 80_=1.364, p=0.2539).

**Figure 4.**
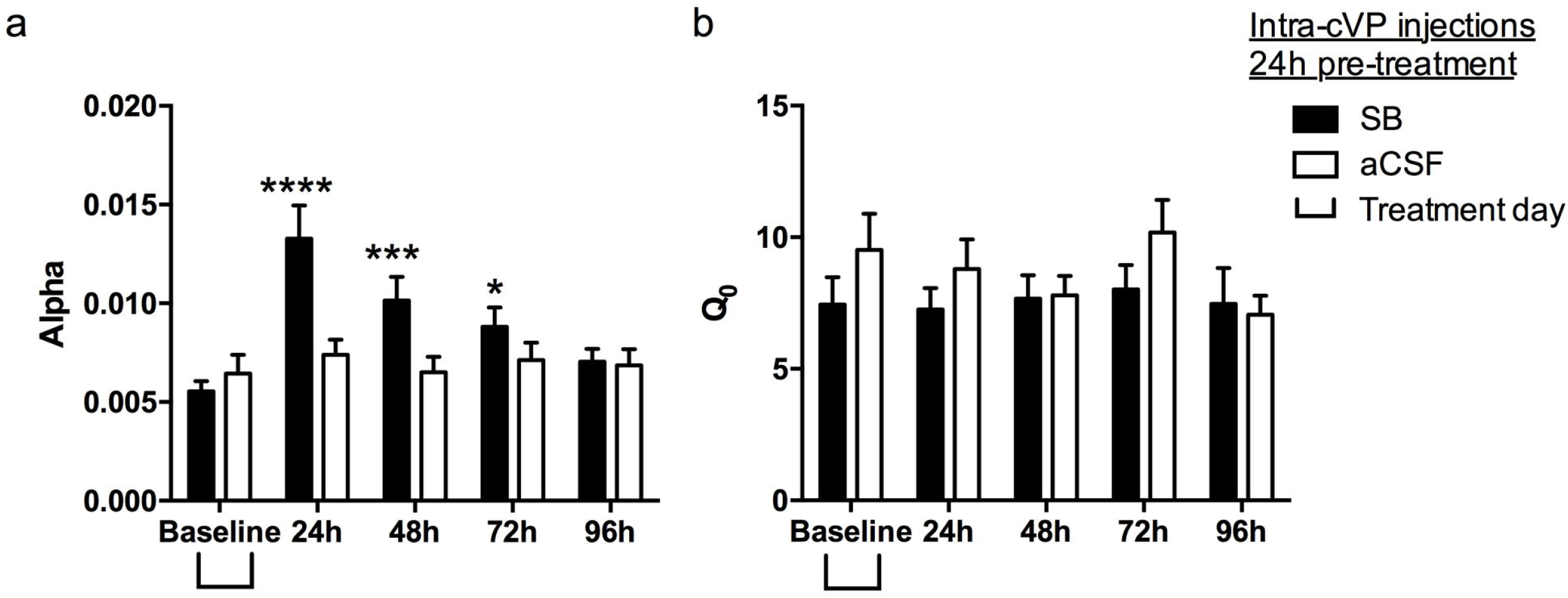
Intra-cVP pretreatment with SB reduced motivation for remifentanil for up to 72h. **(a)** Following BE testing at 24h post intra-cVP SB or aCSF treatment (presented in Figure 3), rats were tested (with no further SB administration) on the BE paradigm for an additional 3 sessions. SB pretreatment was associated with a significant increase in *α* values (decreased motivation) up to 72h post-injection. There was no effect of vehicle treatment. **(b)** Pretreatment with SB or aCSF had no effect on remifentanil consumption under null cost conditions at any time point. Bar graphs represent mean ± standard error of mean (SEM). *p<0.05, ***p<0.001, ****p<0.0001, compared with baseline values.

#### 2.3.2. Intra-cVP pretreatment with SB (24h prior to testing) attenuated cued reinstatement of remifentanil seeking

To test whether intra-cVP pre-treatment with SB might also be effective at blocking cued reinstatement at a later time point, following BE testing we extinguished lever pressing for remifentanil over a period of 7d (**Fig. 5a**). Animals were then microinjected with either SB or aCSF (counterbalanced) bilaterally into cVP, 24h prior to cued reinstatement test sessions. A 2×3 factor repeated measures ANOVA revealed a significant main effect of ‘Treatment’ (F_2,32_=7.765, p=0.0018) and ‘Lever’ (F_2,32_=18.82, p=0.0005), and a significant ‘Treatment’ × ‘Lever’ interaction (F_2,32_=8.583, p=0.0010). Dunnett post-hoc analyses showed that remifentanil-associated cues reinstated active lever pressing after vehicle pretreatment (vehicle vs. extinction, p=0.0010; **Fig. 5b**). SB in cVP significantly reduced responding on the active lever during reinstatement (SB vs. vehicle, p=0.0359; **Fig. 5b**). This reinstatement was blocked by pretreatment with SB into cVP (p=0.4289 versus extinction levels). There was no change in responding on the inactive lever (**Fig. 5b**; F_2, 26_=0.2993, p=0.6876) after pretreatment with SB (p=0.91) or vehicle (p=0.60).

**Figure 5.**
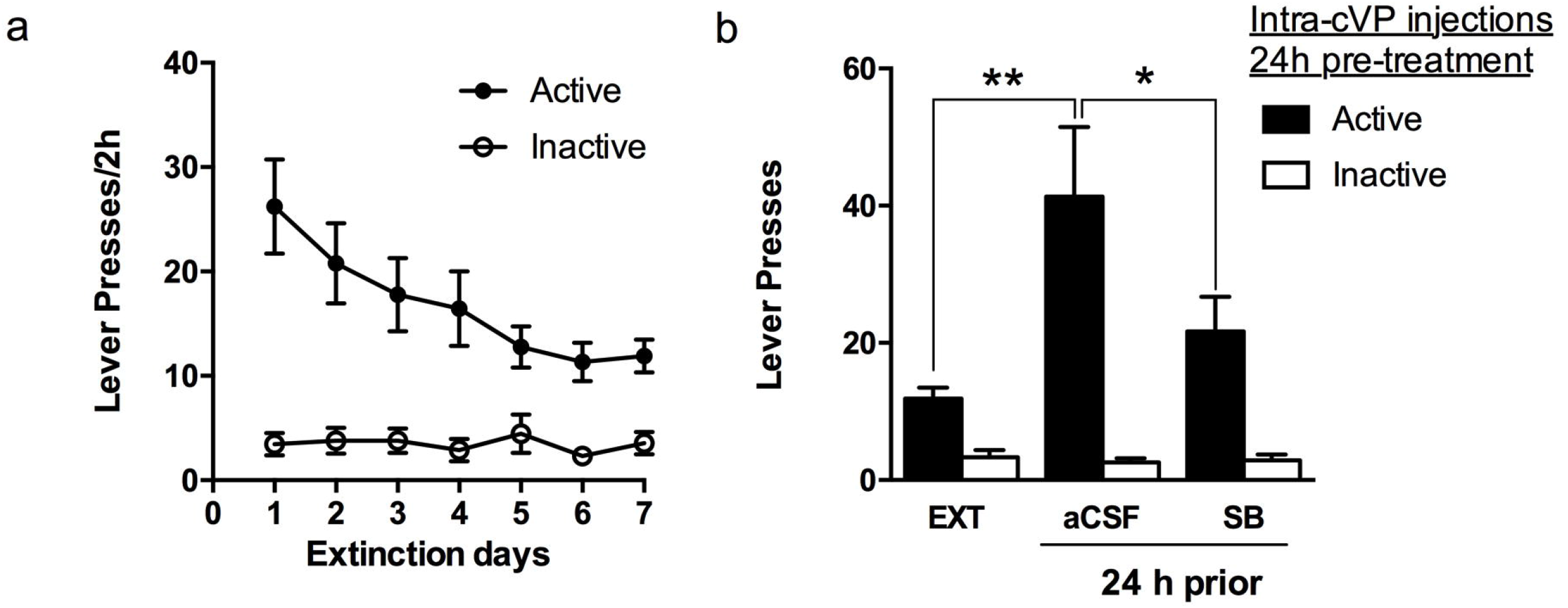
Intra-cVP pretreatment with SB 24h prior to testing reduced cued reinstatement of remifentanil seeking. **(a)** Following BE testing, lever pressing was extinguished in daily sessions for 7d. Data depict the mean number of active and inactive lever presses over these 7d. **(b)** Intra-cVP pretreatment with SB 24h prior to testing was associated with a significant decrease in the number of active lever responses during a 2h cued reinstatement test compared to aCSF treatment. There was no effect of treatment on inactive lever responding. Bar graphs represent mean ± standard error of mean (SEM). *p<0.05, **p<0.01. n=9 for all groups.

#### 2.3.3. No effect of intra-cVP pretreatment with SB on general locomotor activity

To test for non-specific motoric effects of SB pretreatment, rats were tested for locomotor activity 24h following intra-cVP pretreatment with SB or vehicle. Pretreatment with SB had no effect on the total distance traveled (**Fig. 6a**; F_23,368_=1.467, p=0.0779), nor was there an effect on either horizontal (**Fig. 6b**; p=0.202) or vertical (**Fig. 6c**; p=0.395) activity over the 2h test sessions, indicating that was no overall effect of pretreatment with SB on general motor behavior.

**Fig 6.**
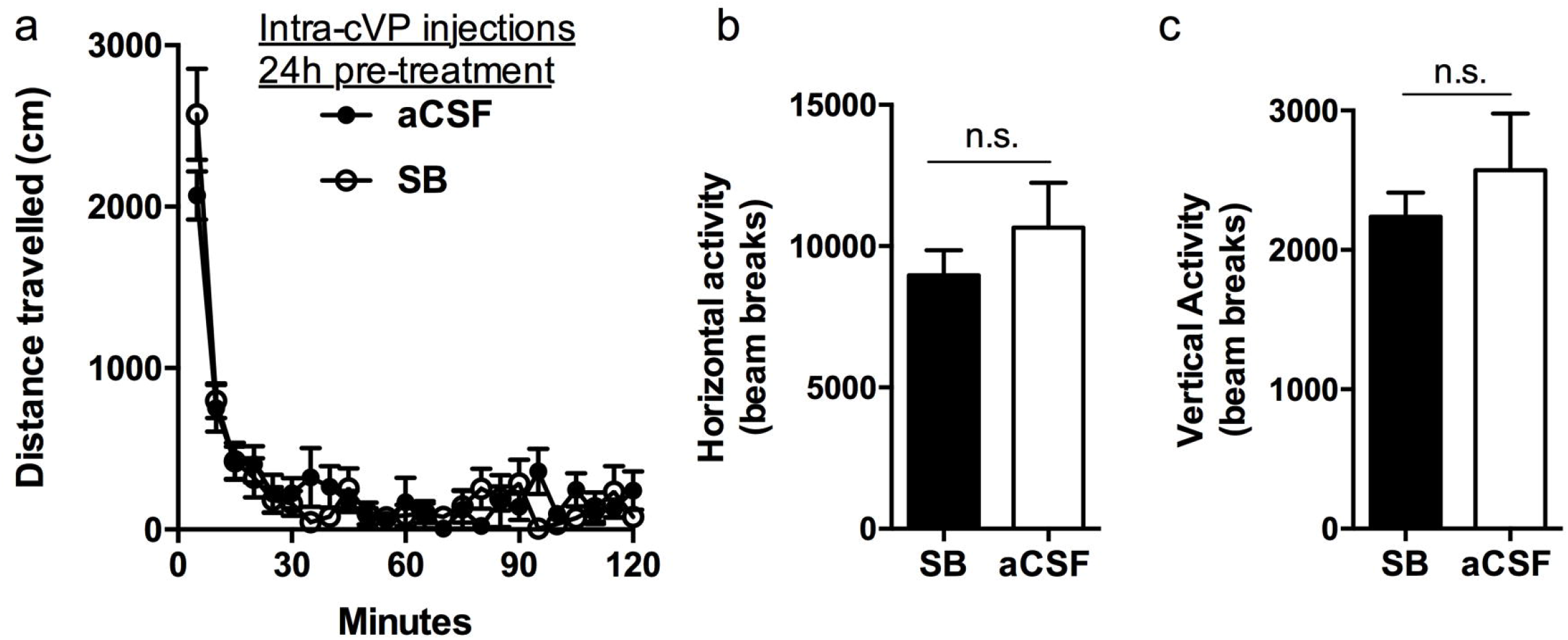
Intra-cVP pretreatment with SB 24h prior to testing had no effect on general locomotor activity. **(a)** Rats displayed no differences in total distance traveled in a test of general locomotor activity when treated with intra-cVP SB 24h prior to testing. **(b,c)** SB treatment also had no effect on horizontal (b) or vertical (c) activity. Bar graphs represent mean ± standard error of mean (SEM). n.s.: not significantly different. n=9 for all groups.

## 3. Discussion

We tested whether inhibition of remifentanil-seeking by the selective Ox1R antagonist SB persisted beyond the pharmacological availability of the drug. We found that when administered systemically, SB reduced motivation (decreased economic demand) for remifentanil for up to 24h post-treatment (Experiment 1). When administered locally into cVP, the anti-motivational effects of SB persisted for at least 48h (Experiment 2). In a separate group of rats that were treated with intra-VP SB 24h prior to behavioral testing, we observed reduced motivation for remifentanil in 3 subsequent behavioral tests, indicating that the effects persisted for 72h post-SB administration. We also showed that cued reinstatement of remifentanil seeking was decreased 24 hours after intra-cVP SB (24h prior), without affecting general locomotor activity. Together, these results indicate a persistent effect of SB, both when delivered systemically or intra-VP, on motivated responding for the opioid remifentanil.

We observed reduced motivation for remifentanil at 24h, but not 48h, following systemic SB treatment. This finding aligns with those of Brodnik et al. (2018) who recently tested the persistence of SB effects on cocaine self-administration on a discrete trials-3 schedule of reinforcement. In this paradigm, rats are allowed access to cocaine in 10min trial blocks that are initiated 3 times each hour over each 24h period. This schedule promotes a cyclic pattern of drug taking across the day characterized by high levels of drug intake during the active (lights off) period and low intake during the inactive (lights on) period. The authors reported that when given systemically during the active (lights off) period, SB accelerated the termination of drug taking *and* delayed the re-initiation of drug taking in the following active period (15-21h following SB delivery). Our data are also consistent with several previous studies which reported that acute treatment with SB (30mg/kg, i.p.) suppressed intake of home cage chow and high fat food at 24h post-dosing, an effect that was also accompanied by a significant loss of body weight over this period (Haynes et al., 2000; Ishii et al., 2005; White et al., 2005). Normal feeding behavior was restored at 48h post-SB treatment (Ishi et al, 2005), which is consistent with the time we observed a normalization of remifentanil-seeking behavior (although body weight deficits are maintained for up to 3d post treatment (Rodgers et al., 2001; White et al., 2005). Together, these findings point to a common time course of SB’s efficacy following systemic administration across several reinforcers, including psychostimulants, opioids and food.

The pharmacokinetics of systemic SB treatment have been well-characterized in adult male rats. At 30mg/kg (i.p.), SB reaches peak plasma and brain concentrations at ~30 min and has a half-life of ~4h (Ishii et al., 2005). At 10mg/kg (i.p.; a dose also commonly used in tests of drug motivation), SB has a terminal elimination half-life of ~0.4h (Porter et al., 2001). As the compound is virtually undetectable in plasma or brain at 2h and 8h post-dosing (10 and 30mg/kg dose, i.p., respectively) (Porter *et al*., 2001; Ishii *et al*., 2005), prolonged occupancy at the Ox1R would seem unlikely as the potential mechanism underlying the persistent effects reported here. One hypothesis is that blockade of Ox1R produces enduring neuroadaptations to the Ox1R or downstream signaling pathways involved in motivation (Ishii et al., 2005). Indeed, Brodnik et al. (2018) reported that intra-VTA administration of an alternative Ox1R antagonist, RTIOX-276 (RTI) delivered 24h prior to testing produced a robust reduction in accumbal DA uptake inhibition following a cocaine challenge. These authors also reported a significant reduction in accumbal dopamine transporter (DAT) phosphorylation at 24h post-RTI administration. It is unclear whether such changes could underlie the effects observed here, especially given that accumbal dopamine signaling may be less critical for opioid reward behavior compared to psychostimulants (Gerrits et al., 1994; Gerrits and Van Ree, 1996; Pettit et al., 1984), although see (Corre et al., 2018). Thus, it will be critical for future studies to examine how orexin peptides acting at Ox1R is important for the normal functioning of opioid reward pathways

Many studies utilize local microinjections of SB to examine the role of orexin signaling in specific reward loci. VP is a well-characterized reward structure with major reciprocal connections to the mesolimbic dopamine system, including the nucleus accumbens (NAc) and ventral tegmental area (VTA) (Groenewegen et al., 1993; Haber et al., 1985). Accordingly, VP is a critical node in the brain reward circuit that underlies a range of drug-seeking behaviors, including self-administration and reinstatement behavior (Churchill and Kalivas, 1994; Cooper et al., 2017; Heinsbroek et al., 2017; Mahler et al., 2014b; Smith and Berridge, 2007). VP (particularly cVP) receives dense innervation from orexin containing neurons in LH, and both Ox1R and Ox2R are expressed in this region. Ho and Berridge provided the first evidence that orexin acts at cVP to mediate reward by showing that intra-cVP infusions of orexin-A enhance the hedonic properties of sucrose (Ho and Berridge, 2013). We recently extended these findings to show that blockade of Ox1R signaling in cVP reduces motivation and low-effort remifentanil consumption on an economic demand task that commenced immediately following SB injections (Mohammdkhani et al., 2018). Here, we show that these effects persist for at least 48h following treatment. These effects were not attributable to reduced remifentanil intake on the initial BE test, as similar effects were observed following SB pretreatment 24h prior to testing (reduced demand was observed up to 72h under these conditions). Pretreatment with SB 24h prior to testing was also associated with reduced cued reinstatement behavior, but no change in general locomotor activity. Together these findings indicate that the effects of SB on motivated behavior are more persistent following intraparenchymal microinjections compared to systemic injections.

To our knowledge, this is the first study to show a persistent effect of local SB microinjections on reward seeking behavior. Unlike systemic dosing, little is known about the time course of SB availability following intraparenchymal injections, nor is it clear whether this type of administration is associated with persistent neuroadaptations at the site of injection, and how these might relate to reward seeking. VP receives substantial dopaminergic input arising from VTA (Klitenick et al., 1992; Root et al., 2015; Smith and Kieval, 2000), and VTA inputs to VP are activated by drug cues (Mahler and Aston-Jones, 2012). We observed effects of SB up to 72h post-treatment following local injections (compared to 24h post-systemic treatment); this might be related to a higher concentration of SB in the brain following the intraparenchymal route of delivery. Future studies should also seek to examine whether similar persistent effects are observed following local SB injections into other regions where Ox1R signaling is known to regulate drug behavior, including VTA (James et al., 2011; Mahler et al., 2013; Wang et al., 2009).

Our findings have important implications with respect to the design of studies involving Ox1R antagonists to test the role of Ox1R signaling in reward seeking paradigms. Indeed, care should be taken when interpreting behavioral results from testing conducted between 24h (for systemic) and 72h (for intracranial) post-administration, as behavior is likely to be influenced by the prior treatment at these time points. Moreover, while only relatively few studies have examined the effect of repeated/chronic SB dosing (Borgland et al., 2006; Zhou et al., 2012), such studies will be important for understanding the potential clinical use of Ox1R antagonists for the treatment of disorders such as addiction. These studies should seek to understand whether the persistent effects, and possible neuroadaptations, associated with acute dosing reported here are additive with repeated dosing.

## 4. Materials and methods

### 4.1. Subjects

A total of 35 male Sprague Dawley rats (initial weight 275–300 g; Charles River, Raleigh, NC, USA) were used in these experiments. Data from 24 of these rats were analyzed for other purposes in a separate publication (Mohammadkhani et al., 2018); the remaining 11 rats were prepared exclusively for this study. Rats were single-housed and maintained under a 12 h reverse light/dark cycle (lights off at 08:00 h) in a temperature and humidity-controlled animal facility at Rutgers University. Food and water were available ad libitum. All experimental procedures were approved by the Rutgers Institutional Animal Care and Use Committee and were conducted according to the Guide for the Care and Use of Laboratory Animals. Rats were handled daily after a 3-day acclimation period at the facility; all experiments were performed in the rats’ active (dark) phase.

### 4.2. Drugs

Remifentanil (obtained from the NIDA Drug Supply Program, National Institute of Drug Abuse) was dissolved in 0.9% sterile saline for intravenous (iv) self-administration. SB334867 (SB), a selective antagonist for the Ox1R (supplied by the NIDA Drug Supply Program, National Institute of Drug Abuse) was dissolved in sterile artificial cerebrospinal fluid (aCSF) at a concentration of 1mM for cVP microinjections. For systemic administration, SB was suspended in 10% 2-hydroxypropyl-b-cyclodextrin in sterile water and 2% dimethyl sulfoxide (Sigma-Aldrich, St. Louis, MO, USA). Three different group of rats were treated with SB as follows: In experiment 1, rats were injected with 30 mg/kg SB or vehicle (i.p.) at a volume of 4.0 ml/kg, 30 min prior to the first behavioral test (n=10). Rats in Experiment 2 received an intra-cVP microinjection of SB (0.3 μl) immediately prior to the first behavioral test (n=14). In Experiment 3 rats received intra-cVP microinjection of SB (0.3 μl) 24h prior to the first behavioral test (n=11). A within-subjects design was used whereby each rat received both SB and vehicle; the order was counterbalanced across subjects.

### 4.3. Intravenous catheter surgery

Following acclimation to the animal facility, rats were anesthetized with a ketamine/xylazine mixture (56.5 and 8.7 mg/kg, i.p., respectively) followed by an injection of an analgesic (rimadyl, 5 mg/kg; s.c.). Rats were then implanted with an indwelling catheter (modified 22 ga cannula, Plastics One) into the jugular vein that exited the body via a biopsy on the back caudal to the mid-scapular region. Catheters were flushed with cefazolin (0.1 ml; 100 mg/ml) and heparin (0.1 ml; 100 U/ml) immediately following surgery, daily beginning 3 days after surgery, and continuing throughout self-administration. Rats were allowed to recover for a minimum of 7 days after surgery before remifentanil self-administration training.

### 4.4. Stereotaxic surgery

Immediately following catheter implantation, animals in Experiments 2 and 3 were placed in a stereotaxic frame (Kopf, Tujunga, CA, USA) and implanted with bilateral stainless steel guide cannulae (22 gauge, 11 mm, Plastics One, Roanoke, VA, USA) 2mm dorsal to VP (coordinates relative to bregma skull surface in mm: −0.8 posterior, ±2.6 medial–lateral, −7.5 ventral; (Paxinos and Watson, 1998). The cannulae were secured to the skull using jeweler’s screws and dental acrylic; stylets were placed into the guide cannula to prevent occlusion.

### 4.5. Self-administration training

The self-administration procedure was similar to a recently published study from our laboratory (Porter-Stransky et al., 2017). All self-administration sessions occurred in operant chambers controlled by Med-PC IV software (Med Associates). Each operant chamber was equipped with two levers, a cue light above each lever, a tone generator, a house light, and an infusion pump. A timeline of experimental training and tests is found in Fig. 3a. A response on the active lever resulted in remifentanil infusion (1 μg delivered over 4 sec) paired with a discrete compound cue (auditory tone plus white cue light above the active lever) during self-administration. After each infusion, a 20-sec time-out occurred (signaled by turning off the house light) when additional presses were recorded but did not yield remifentanil or cues. Presses on the inactive lever were recorded but had no consequences. Each session ended after 2h or when 80 infusions were earned, whichever first occurred. Subjects were trained on the FR1 procedure for a minimum of 6 sessions. After earning a minimum of 25 infusions within 2h for at least 3 consecutive sessions, animals advanced to the behavioral economics threshold procedure. Data for self-administration training and BE testing for Experiments 1 and 2 are reported elsewhere (Mohammdkhani et al., 2018).

### 4.6. Behavioral-economics threshold procedure

Economic demand training for remifentanil was conducted as in a recent study from our laboratory (Porter-Stransky et al., 2017). As previously described (Bentzley et al., 2013), the BE procedure varies the cost of drug by changing the amount of drug infused while maintaining an FR1 schedule of reinforcement. The dose of remifentanil resulting from a response on the active lever decreased every 10 minutes; therefore, each 110-minute session tested 11 doses of remifentanil (2, 1, 0.6, 0.3, 0.2, 0.1 0.06, 0.03, 0.02, 0.01 and 0.006 μg/infusion). As in FR1 training, each infusion was accompanied by presentation of a light-tone compound cue; responses on the inactive lever had no consequence. However, unlike the FR1 training procedure, there were no limits on how much drug rats could take in a session nor were there time-out periods in which drug was unavailable. Subjects were trained daily on the BE procedure until responding became stable, defined as ≤20 percent variation in α and Q_0_ for at least 3 consecutive days. Rats then began treatment, as outlined below.

### 4.7. Extinction and reinstatement testing

Rats in Experiment 3 underwent extinction training in daily 2h sessions, where lever presses yielded neither remifentanil nor cues. Rats received extinction training for a minimum of 7 days and until they met the criteria of ≤15 active lever presses for at least two consecutive sessions. The following day, rats were tested for cued reinstatement of remifentanil seeking; responses on the active lever during these sessions resulted in presentation of the cues previously paired with remifentanil infusions, but no remifentanil. Rats received at least 2 additional days of extinction (≤15 active lever presses criterion) before undergoing an additional reinstatement test (James et al., 2018b; McGlinchey et al., 2016). The order in which rats received treatment was counterbalanced.

### 4.8. Locomotor testing

Rats in Experiment 3 were also tested for general locomotor activity, as described previously (James et al., 2018b; McGlinchey et al., 2016). Rats were placed in locomotor chambers (clear acrylic, 42 × 42 × 30 cm) equipped with SuperFlex monitors (Omnitech Electronics Inc, Columbus, OH) containing a 16 × 16 photobeam array for the x/y-axis (horizontal activity) and 16 photobeams for the z-axis (vertical activity). Photobeam breaks were recorded by Fusion SuperFlex software. Rats were given one 2h session to habituate to the chamber. On test days, rats received intra-cVP injections of SB or vehicle immediately before being monitored for locomotion over a 2h period; total distance traveled, as well as horizontal and vertical activity were recorded. Rats underwent two tests (drug and vehicle treatments), and the order of treatments was counterbalanced.

### 4.9. Experimental design

#### 4.9.1. Experiment 1: Systemic SB30 administration immediately prior to initial behavioral economic testing for remifentanil

Rats were trained to self-administer remifentanil and were tested on the BE paradigm until stable α and Q_0_ values were obtained. The following day, rats were treated systemically with SB30 or vehicle and tested on the BE paradigm 30mins later (0.5h). Rats were re-tested on the BE paradigm in a drug-free state on two additional sessions, 24h apart (24h and 48h posttreatment). Rats were returned to their home cages between testing sessions. Each animal continued on daily BE sessions until α and Q_0_ values were within ±20% of pre-treatment values (baseline); rats then underwent testing as before with the opposite treatment (SB or vehicle; order counterbalanced). Full analyses of data from the 0.5 h test is presented elsewhere (Mohammdkhani et al., 2018) and data presented here represent a re-analysis of behavioral testing on this initial test and during washout days.

#### 4.9.2. Experiment 2: Intra-cVP microinfusions of SB immediately prior to testing for remifentanil economic demand

Local microinfusions of SB are often used to identify important sites of orexin signaling in reward seeking. Such studies have reported that orexin acts at cVP to mediate motivational and hedonic properties of reward (Ho and Berridge, 2013; Mohammdkhani et al., 2018); however, these effects have only been tested acutely. Here we sought to test the persistence of these effects by testing rats on the BE task at 0h following an intra-cVP microinjection of SB, and then re-testing in a drug-free state at 24h and 48h. Each rat received only a single injection of SB into cVP.

The design of this study was identical to that of Experiment 1, except cVP microinjections replaced systemic SB injections. On the day prior to testing, rats were acclimated to the infusion procedure by inserting injectors (28g) bilaterally into the guide cannula that protruded 2.0mm below the bottom of the cannulae; injectors were left in place for 1 min (no infusions were made). Immediately prior to testing on the following day, rats received bilateral microinfusions (0.3μl/side) of either SB (1mM) or aCSF. The order of SB and vehicle microinfusions were counterbalanced and administered via polyethylene tubing connected to gastight 10-μl Hamilton syringes (Hamilton, Reno, NV, USA) set in an infusion pump (Model 975, Harvard Apparatus, Holliston, MA, USA).

We also performed control microinfusions of SB to confirm that any behavioral effects were not due to actions at a dorsal site because of diffusion of SB along the cannula tract. Injections were made dorsal to VP using injectors that projected only 0.2 mm below the tip of the guide cannulae. Dorsal control microinjections were performed in a session prior to cVP injection sessions. Control microinjections were made immediately prior to the first BE test (0h).

Similar to *Experiment 1*, full analyses of data from the 0h test is presented elsewhere (Mohammdkhani et al., 2018) and data presented here represent a re-analysis of behavioral testing on this initial test and during washout days.

#### 4.9.3. Experiment 3: Pretreatment with intra-VP SB 24h prior to initial behavioral economic and cued-reinstatement testing

It is possible that reduced remifentanil intake on the initial BE test following acute SB administration (i.e. at 0h) could affect performance on subsequent tests (i.e. at 24 and 48h). To address this, we prepared a new group of animals with cVP cannulae and administered intra-cVP microinfusions of SB or aCSF, and then underwent BE testing at 24h, 48h, 72h, and 96h post-treatment. Once BE values returned to pre-treatment values, rats were tested under the opposite treatment condition (aCSF vs SB).

Because this is the first study (to our knowledge) to test the effect of pre-treating with SB locally 24h prior to behavioral testing, we also examined the effects on cued reinstatement of remifentanil seeking, as this behavior is well-documented to be orexin-dependent. Remifentanil-seeking was extinguished and rats were tested for cued reinstatement behavior (as described above). Intra-cVP microinfusions of SB or aCSF (counterbalanced) were delivered 24h prior to reinstatement testing (following the final extinction session).

Rats were also assessed for general locomotor activity; rats received intra-cVP microinjections of SB or aCSF 24h prior to motor assessment (described above). The order in which rats were tested on reinstatement and locomotor tests after BE was fully counterbalanced to account for any order effects associated with repeated intra-cVP microinjections.

### 4.10. Localization of injection sites

After the final behavioral test, rats in *Experiments 2* and *3* were deeply anesthetized with ketamine/xylazine (56.6/8.7mg/kg) and received bilateral microinfusions of pontamine sky blue (0.3μl) to mark the locations of the injectors. Rats were then decapitated, and brains were flash-frozen in 2-methylbutane and stored at – 80°C. Brains were sectioned into 40 μm-thick sections on a cryostat (Leica CM 3050), mounted, Nissl-stained with neutral red, and cover slipped to localize cannula tracts and verify injection sites. A total of 5 rats received misplaced injections that were instead directed at the rostral VP (rVP) – an analysis of the acute effects of SB in these animals is presented elsewhere (Mohammdkhani et al., 2018). Here, we focused our analyses on all other animals that received accurate SB injections directed at cVP.

### 4.11. Data analysis

Demand curves (example presented in Fig.3d) were generated for each BE session as previously described (Bentzley et al., 2013; Porter-Stransky et al., 2017). An exponential demand equation (Hursh and Silberberg, 2008) was applied to each animal’s data to generate a demand curve and estimates of preferred remifentanil intake at zero cost (Q_0_, where the computed demand curve intercepted the ordinate) and demand elasticity (α, the slope of the demand curve). Larger α values indicate greater demand elasticity and are characterized by a greater reduction in responding as drug price increases; this is interpreted as decreased motivation as described in our previous work (Bentzley et al., 2013). Smaller α values indicated less demand elasticity and indicate continued responding for drug despite increases in the cost to obtain drug (i.e., increased motivation). Curve fitting was performed similarly to our previous studies, whereby all data points up until two bins past the point at which maximal responding was observed (O_max_) were included in the generation of demand curves. Unlike our previous cocaine studies where data from the first ‘load up’ bin were excluded from analyses, this was unnecessary here because subjects were not observed to ‘load up’ on remifentanil, as has been reported elsewhere (Mohammdkhani et al., 2018; Panlilio et al., 2003; Porter-Stransky et al., 2017).

Parametric and nonparametric statistical analyses were performed in Graphpad Prism version 6. Acquisition of self-administration, individual differences in BE parameters and the effects of SB on locomotor activity were analyzed using repeated measures ANOVA (total distance traveled) and paired t-tests (horizontal/vertical activity). The effects of SB on active/inactive lever responding during reinstatement tests was assessed using separate twoway repeated measures ANOVAs with ‘treatment’ (vehicle, SB) and ‘lever type’ (active, inactive) as the variables. Corrections (Dunnett) were applied to post hoc tests to reduce the risk of Type 1 errors. Linear regression was used to correlate individual Q_0_ and α values. Data for self-administration training and BE testing for Experiments 1 and 2 are reported elsewhere (Mohammdkhani et al., 2018).

## Acknowledgements

We would like to thank Drs. Kirsten Porter-Stranksy and Brandon Bentzley for their assistance with the remifentanil demand protocol. We would also like to thank Dr. Hannah Bowrey and Caroline Pantazis for their helpful guidance in interpreting the data, as well as Nupur Jain for her assistance.

